# Persistence of phenotypic responses to short-term heat stress in the tabletop coral *Acropora hyacinthus*

**DOI:** 10.1101/2022.05.18.492444

**Authors:** Nia S. Walker, Brendan H. Cornwell, Victor Nestor, Katrina C. Armstrong, Yimnang Golbuu, Stephen R. Palumbi

## Abstract

Widespread mapping of coral thermal resilience is essential for developing effective management strategies and requires replicable and rapid multi-location assays of heat resistance and recovery. One- or two-day short-term heat stress experiments have been previously employed to assess heat resistance, followed by single assays of bleaching condition. We tested the reliability of short-term heat stress resistance, and linked resistance and recovery assays, by monitoring the post short-term heat stress phenotypic response of fragments from 101 *Acropora hyacinthus* colonies located in Palau (Micronesia). Following short-term heat stress, bleaching and mortality were recorded after 16 hours, daily for seven days, and after months one and two into recovery. To follow corals over time, we utilized a qualitative, non-destructive visual bleaching score metric that correlated with standard symbiont retention assays. The bleaching state of coral nubbins 16 hours post-heat stress was highly indicative of their state over the next 7 days, suggesting that symbiont population sizes within corals may quickly stabilize post-heat stress. Bleaching 16 hours post-heat stress predicted likelihood of mortality over the subsequent 3-5 days, after which there was little additional mortality. Together, bleaching and mortality suggested that rapid assays of the phenotypic response following short-term heat stress were good metrics of the total heat treatment effect. Additionally, our data confirm geographic patterns of intraspecific variation in Palau and show that bleaching severity among colonies was highly correlated with mortality over the first week post-stress. We found high survival (98%) and visible recoverability (100%) two months after heat stress among coral nubbins that survived the first week post-stress. These findings help simplify rapid, widespread surveys of heat sensitivity in *Acropora hyacinthus* by showing that standardized short-term experiments can be confidently assayed after 16 hours, and that bleaching sensitivity may be linked to subsequent survival using experimental assessments.

## Introduction

There is urgent need for research that aims to uncover the mechanisms leading to coral stress resilience (Hughes et al., 2003; Hughes et al., 2017), which is the ability of these keystone organisms to survive variable and hostile environments. Whether corals are resilient in the face of mounting environmental challenges depends on a number of biotic and abiotic factors, including respective tolerance limits of corals and dinoflagellate endosymbionts and coral host-symbiont interactions (Rowan 2004; Howells et al., 2012; Bay et al., 2016; Drury 2020), coral species composition (Berumen and Pratchett 2006; Alvarez-Filip et al., 2013), nutrient availability (Morris et al., 2019), and the duration and intensity of stressors (Oliver and Palumbi 2011; Couch et al., 2017; Klepac and Barshis 2020).

The importance of resilience to future reef function has led to calls for increased attention toward mapping heat resistance across and within species (Cornwell et al., 2021). To accommodate the large number of experiments needed for such mapping, researchers have recently focused on experiments that impose short pulses of heat exposure mimicking high temperatures in shallow waters at noon time low tides (Ruiz-Jones and Palumbi, 2017; Grottoli et al., 2021). These heat pulse experiments impose high heat for short periods and then typically assay coral bleaching the morning after (see Grottoli et al., 2021). Based on these experiments, immediate impacts of heat stress on corals can be observed in the transcriptome, including upregulation of genes associated with the immune response and apoptosis (Barshis et al., 2013; Seneca and Palumbi, 2015; Louis et al., 2017; Traylor-Knowles et al., 2017). Symbiont impacts, for example, cellular structure damage and oxidative stress during and hours after exposure to stressors may also influence coral holobiont survival (Hoogenboom et al., 2012; Downs et al., 2013; Oakley et al., 2022; Zhang et al., 2022).

Coral heat resistance studies are primarily concluded within the first 16 hours post-heat stress (McLachlan et al. 2020), and there is comparatively little information about heat stress impacts beyond the first 24 hours and over the first few days after a heating event. Better understanding heat stress impacts beyond the immediate effects and hours afterwards would further illuminate how the coral holobiont manages stress and then transitions from a stressed state to a recovery mindset. This may provide further validation for the utility of such short-term experiments and rapid assays in the extensive reef mapping projects that may be necessary to find, protect, and manage future reefs. Extending the timeline of heat stress experiment observation could also result in directly linking coral heat stress resistance and recovery, which may be especially important when there is high variation in resistance and recovery ability within and between species.

In this study we investigated early phenotypic responses to heat stress in coral nubbins (i.e., fragments) collected from individual colonies of *Acropora hyacinthus*—an abundant reef-building coral species with extensive geographic distribution that is representative of many widespread, bleaching sensitive taxa inhabiting tropical reefs (Marshall and Baird 2000; Strychar et al., 2004; Sakai et al., 2019; Cornwell et al., 2021). We employed a two-day short-term heat stress experiment followed by daily monitoring of bleaching intensity and mortality for seven days, to test the stability of the bleaching response and whether bleaching severity is linked to higher likelihood of mortality shortly after heat stress. We additionally revisited the experimental system approximately one and two months after heat stress to determine how variation in heat stress resistance and short-term impacts of the heat stress response affected longer-term recovery.

To allow for repeated and non-destructive bleaching measurements, we employed a five-point visual bleaching score (VBS) system based on coloration relative to baseline coral nubbin color throughout the post-heat stress period. This qualitative bleaching method allowed for simple, undisturbed observation of sample bleaching state following heat stress. As with other studies that used visual bleaching metrics (Thomas and Palumbi 2017; Manzello et al., 2019; Morikawa and Palumbi 2019; Thomas et al., 2019; Fuller et al., 2020; Ritson-Williams and Gates 2020), we corroborated results with a quantitative symbiont density metric. Therefore, our daily observational study was able to provide further insight into the heat stress response by examining this short timeframe following heat stress. This study highlights the importance of combined coral heat stress resistance and recovery assays and considering survival and recovery on short timescales.

## Materials and Methods

### Coral Colony Sampling

We sampled 101 *Acropora hyacinthus* colonies located on 28 reef locations in Palau’s northern and southern lagoons (Fig. S1). We selected on average 3-4 colonies per reef, a subset of colonies from a larger coral reef survey program (Cornwell et al., 2021). Sampling took place from 21^st^ July to 8^th^ August 2018 (Table S1). The heat stress experiments were conducted on an ongoing basis, i.e., coral nubbins were added to tanks as they were collected over staggered days. The primary intention of widespread sampling was to capture heat resistance variability among a set of previously sampled coral colonies with known resistance history (Cornwell et al., 2021).

Temperature on reefs was recorded from 8^th^ November 2017 to 8^th^ August 2018 at 10 min intervals (HOBO, OnSet Computing, Massachusetts). Loggers were placed adjacent to a subset of colonies on reefs. Temperature data were averaged for reefs with multiple retrieved loggers; replicate loggers were analogous within reefs. Loggers were also irretrievable from 8 out of 28 reef sites (Table S1). To quantify temperature differences for analysis, we counted the number of events above 31°C—this threshold allowed us to widely compare temperature spikes across reefs, and other thermal spike thresholds yielded similar results (e.g., 29°C and 30°C, Palumbi 2021). There were no recorded mass bleaching events in Palau during this collection period, though there were mild levels of accumulated heat stress recorded in 2018 ((NOAA Coral Reef Watch; Skirving et al., 2020). Our data found that reef temperatures had little variability and did not exceed 32.5°C (Palumbi 2021).

We sampled fist sized fragments from colonies, loosely wrapped them in bubble wrap that was previously soaked in seawater (DelBeek 2008), and then transported samples by boat to the Palau International Coral Reef Center (PICRC). We then placed these coral fragments into large flow-through holding tanks and clipped them further into four ∼5cm length nubbins. The following day, we scored all nubbins for bleaching (visual bleaching score, described in the following section) and photographed them prior to beginning the short-term heat stress experiment.

### Metrics for Bleaching Severity and Mortality

To measure corals repeatedly without destructive sampling we used a visual bleaching score (VBS) method based on a five-point scale: (1) no bleaching, (2) slightly discolored, with a small amount of visible bleaching, (3) moderately discolored, clearly bleaching, (4) severely discolored, nearly complete bleaching but with some remaining color, (5) no color, total bleaching (Fig. 1, see also Cornwell et al., 2021). The VBS metric was used before and after the short-term heat stress experiment, daily for one week following bleaching, and when we examined the coral nubbins after approximately one and two months of recovery. We scored coral fragments using two observers for all assessments up to a week post-stress. Only one in-person observer was available for one and two months post-stress timepoints, and these scores were confirmed with photographs. Photographs were taken at the same approximate position and time of day, while monitoring bleaching and mortality. Mortality was determined by examining each nubbin for any presence of tissue on the skeleton. We divided mortality into the following categories: not dead (i.e., none or some visible tissue absence) and dead (i.e., complete absence of tissue).

**Figure 1.**
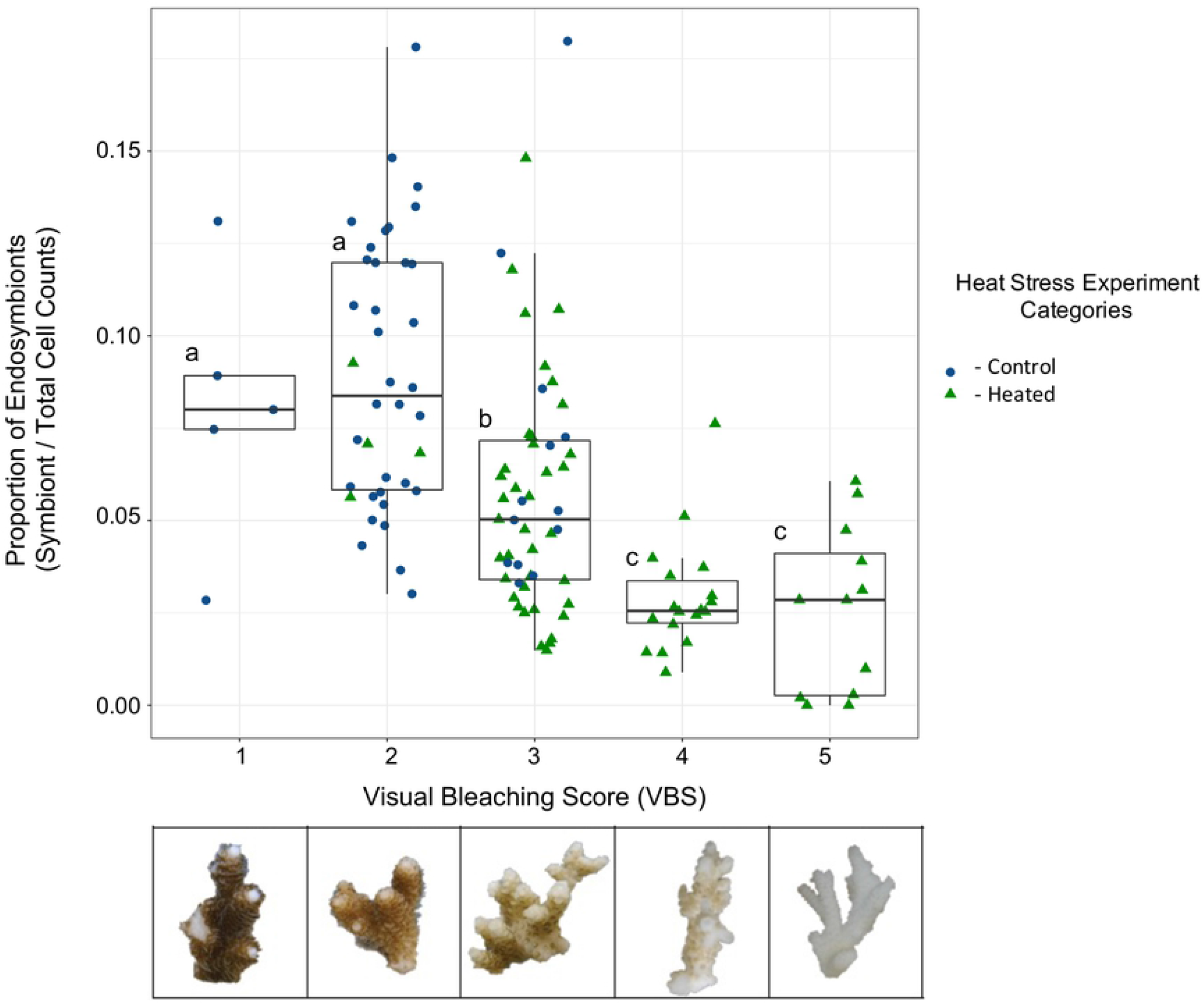
Relationship between symbiont proportion and visual bleaching score metrics after the heat stress experiment. Quantitative measure of bleaching (identified symbiont cells divided by total cell counts) versus qualitative visual bleaching scores (VBS) of sacrificed coral nubbins. Below the x-axis is an example photo representation of the different bleaching severity categories. All sacrificed control (blue circles) and heat stressed (green triangles) nubbins were included to increase sample size across visual bleaching scores. An ANOVA and post-hoc Tukey tests were performed between all visual bleaching score categories, with reef region included as a random intercept to account for spatial variability. We found that categories segregated based on the three labeled groups—a, none to little bleaching; b, moderate bleaching; and c, severe to total bleaching (*p* < 0.05). (Table S2).

Using visual bleaching scores allows for rapid assessment of many corals over consecutive time points and is valuable for long-term assays. However, visual scoring may be subjective among observers or over time, potentially leading to variability or inaccuracy of results. To gauge the value of visual bleaching scores and quantify symbiont concentration after the heat stress experiment, we used flow cytometry (Guava EasyCyte HT; Millepore, Massachusetts) to count the proportion of symbiont cells to total cells in a coral fragment. Immediately after removal from the tanks we airbrushed coral tissue from the skeleton in seawater, centrifuged the slurry and resuspended it in RNAlater. After transport back to Hopkins Marine Station (USA), we washed the RNAlater-tissue suspension in DI water and then resuspended the tissue pellet in a 0.01% SDS:deionized water solution. Samples were homogenized with the PowerGen rotostat for 5 seconds at the highest setting, needle sheared to break apart cell clumps, then diluted by 1:200 in 0.01% SDS and run in triplicate on the flow cytometer. We firstly gated counts on the forward scatter channel (FSC) to exclude small particles (less than 10^2^ fluorescent units). Next, we gated all events that exceeded 10^4^ fluorescent units on the 690 nm detector as symbiont counts (see Krediet et al., 2015). We subtracted events detected in the negative control (0.01% SDS), then we calculated the symbiont proportion as number of symbiont gated events divided by the total number of events after the first FSC gate. We were able to successfully quantify symbiont concentration in 164 nubbins (n=85 controls and n=79 heat stressed samples).

### Short-term Heat Stress Experiment

The short-term heat stress experiments ran in coolers outfitted with seawater inflow (ca. 1/2 volume h^-1^) from the surrounding southern lagoon with large particles filtered out. Tanks were equipped with two chillers (Nova Tec, Maryland), a 300W heater, a submersible water pump (∼280 L h^-1^), and overhanging LED light fixtures (ca. 22-66 μmol m^-2^s^-1^) set to a 12 h light:dark cycle (methods described further in Cornwell et al., 2021). Starting 22^nd^ July 2018, heated coral nubbins ramped from 30°C to 34.5°C over three hours (1000 to 1300 hrs), held at 34.5°C for three hours (1300 to 1600 hrs), then ramped down to 30°C (1600 to 1800 hrs) and held at 30°C until the next day). Two experimental nubbins were subjected to two days of this ramp cycle while two control nubbins sat in separate identical coolers that remained at 30°C for two days per colony. See Cornwell et al., (2021) for evidence of low symbiont concentration variation between two replicate nubbins from these individual colonies, though further sampling may improve characterization of colony heat resistance. The morning after the two-day heat stress experiment, all experimental and control nubbins were scored via visual bleaching score (VBS) and photographed. One replicate experimental and control nubbin per colony were sacrificed in RNAlater for cell counting via flow cytometry. The remaining replicate experimental and control nubbins were placed into a large holding tank for the post-stress experiment period.

### Post-Heat Stress Experiment

Large outdoor flow-through holding tanks with turnover of one full volume per hour were used. Seawater inflow also came from the surrounding southern lagoon, and tank temperature was periodically recorded (∼28.5-30°C). All holding tanks were kept underneath a large roof that provided some light protection, though no additional shade devices were installed to mitigate further light damage, nor were additional light fixtures included. We used water pumps for circulation and relied on natural sunlight. Nubbins were either epoxied upright onto plastic crates or laid flat down on egg crate, and there were no observable differences in survival between methodology (Table S2). Experimental and control nubbins from the same colony were kept next to each other in identical conditions. Corals were added to the holding tank as they finished the heat stress experiment. The morning after the two-day heat ramp was called Day 0 of the post-stress period. At approximately 8:00AM daily until Day 7 post-stress, all coral nubbins were scored for their survival and bleaching severity (via VBS) and were photographed. In order to evaluate visual recovery and mortality approximately one- and two-months post-heat stress, we scored and photographed all corals on: 22^nd^ August, 4^th^ September, 7^th^ September, 10^th^ September, 13^th^ September, and 2^nd^ October. On 10^th^ September 2018 all corals were moved into an adjacent, comparable flow-through holding tank to remove macroalgal buildup on the previous tank’s walls. No macroalgae were removed from the coral nubbins or plastic crate they rested on.

### Statistical Analyses

We ran all statistical analyses in R (version 4.0.5). We tested for differences between heat stressed and control samples’ bleaching severity (Day 0 post-stress, n=88 heat stressed and n=101 control nubbins) and mortality (Day 7 post-stress, n=101 heat stressed and n=101 control nubbins) using Pearson’s chi-square test. We ran a one-way ANOVA and post-hoc Tukey Test to determine whether quantitative symbiont concentration values correlated with qualitative visual bleaching scores. We used a linear mixed effects model to evaluate change in bleaching severity among heat stressed samples from Day 0 to Day 7 of the post-stress period (n=50). We also used a one-way ANOVA and post-hoc Tukey test to evaluate any changes in bleaching on Post-Stress Days 7, 30, and 60 among samples with different levels of heat resistance (based on Day 0 bleaching severity). Further, we ran a mixed effects logistic regression to predict whether bleaching severity on Day 0 influenced likelihood of mortality on Day 60 post-stress (n=88 heat stressed nubbins). We collected coral fragments from a wide geographic range across Palau’s northern and southern lagoons to capture diverse heat stress responses. When evaluating heat resistance and mortality among samples, we accounted for any possible spatial variability between reef regions (see Fig. S1) by using linear mixed effects models (linear regressions and ANOVAs, R package nlme; logistic regression, R package lmer) that specified reef region as a random intercept. Marginal R^2^ (based on fixed effects) and conditional R^2^ (based on fixed + random effects) values were calculated using R package sjstats. Post-hoc Tukey tests for linear mixed effects models were performed using R package multcomp. Reef region had a negligible effect on all statistical analyses (Table S2). Lastly, we also used an ordinal logistic regression to evaluate whether reef temperature might have significantly influenced bleaching severity categories (low: n=13, moderate: n=28, high: n=18 heat stressed nubbins) and mortality (n=8) during the heat stress experiment.

## Results

### Ground truthing VBS with flow cytometry

Symbiont proportion quantitatively tracked visual bleaching scores: low scores (none and visible) were highly distinct from moderate scores and from high scores (severe and total, Fig. 1). In addition, there was little distinction between no bleaching (VBS 1) and visible bleaching (VBS 2), and between severe (VBS 4) and total bleaching (VBS 5) when comparing symbiont proportion averages. On average, corals with little bleaching (VBS 1 and 2) had 8.9 ± 3.6% symbiont proportion, moderately bleached corals (VBS 3) had 5.8 ± 3.4% symbionts, and severely bleached corals (VBS 4 and 5) had 2.8 ± 1.8% symbionts (Fig. 1). These results suggest strong confidence in determining bleaching severity based on visual bleaching scores between minimal and severe bleaching. However, these results also reveal that smaller changes in visible bleaching (i.e., VBS 1 vs 2 and VBS 4 vs 5) as determined with the visual bleaching score method do not correlate well with symbiont proportion. It is worth noting that sample size may have played a role in the ability to accurately capture distinctions between VBS categories—the two categories with the fewest coral nubbins were those with no bleaching (VBS 1, 5 samples) or total bleaching (VBS 5, 12 samples) (Table S1).

### Variable bleaching severity in nubbins immediately following short-term heat stress

After two days of short-term heat stress, 88 of 101 heated branches remained alive but showed a wide variety of bleaching results. Overall, we observed bleaching in 95% of heated nubbins: most nubbins showed visible (17%), moderate (43%) or severe (25%) bleaching (Fig. S2) with one coral showing no bleaching and five being totally bleached. Due to the above quantitative flow cytometry results, corals were partitioned into three bleaching severity categories based on these visual bleaching scores: 22 low bleaching severity (none or visible bleaching), 38 moderate severity (moderate bleaching), and 28 high severity (severe or total bleaching). We excluded the 13 dead coral nubbins from all further bleaching severity analysis.

Controls were similarly evaluated to test for potential negative impacts of sample collection and transport to the laboratory. All control nubbins survived while in the stress tank system, and 96 out of 101 control nubbins showed no or minimal bleaching (Fig. S2). Analysis of control nubbins was particularly important, because samples were collected over a period of 19 days and tested in staggered experiments due to the widespread geography of the reefs sampled. As a result, similar survival and lack of bleaching among controls suggested that experimental artifacts of sampling and handling did not significantly impact observed bleaching severity or mortality in heat stressed nubbins.

### Rapid bleaching results predict bleaching and mortality after one week

After following all bleached corals over the course of one week, we found that regardless of the heat resistance category (low, moderate, or high bleaching) surviving corals had highly stable bleaching severity over time (Fig. 2). The majority of all samples (90%, n = 45 out of 50) remained in the same bleaching severity category from Day 0 to Day 7 post-stress. Of the five corals that switched bleaching severity categories, four improved: 2 moderate ⟶ low bleaching and 2 high ⟶ moderate bleaching. The other coral worsened from moderate to high bleaching. As a result, we found a highly significant positive relationship between visual bleaching scores on Day 0 and Day 7 post-stress (Fig. 3A and Table S2, linear mixed effects model, *p* < 2.2e^-16^, marginal R^2^ = 0.863, df = 44), such that VBS at Day 0 post-stress was highly predictive of VBS throughout the following seven day period.

**Figure 2.**
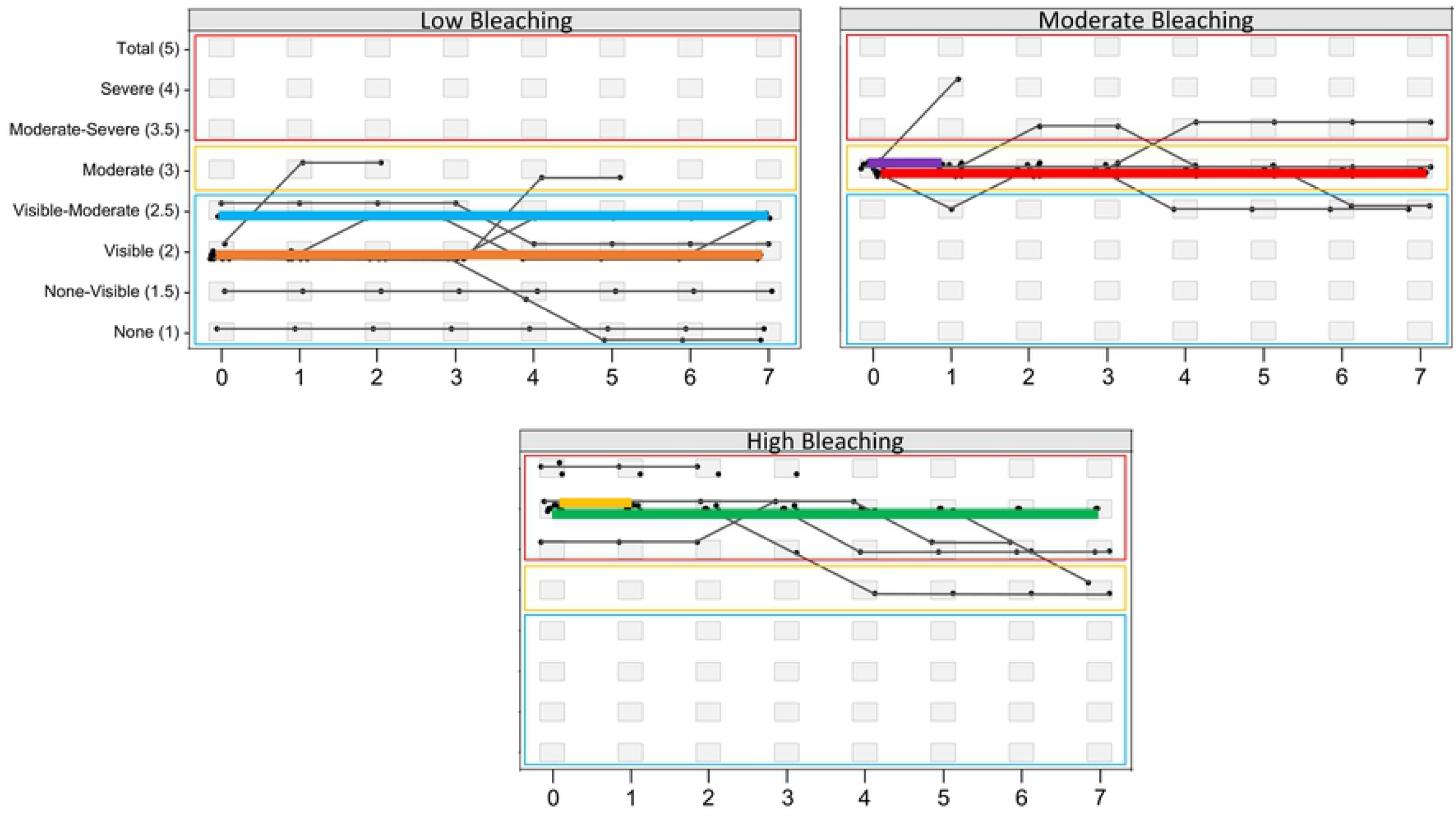
Bleaching stability over one week following heat stress. Ordinal time-series scatterplots, showing bleaching severity and visual bleaching scores of individual bleached coral nubbins daily from Days 0 to 7 of the post-stress period. From left to right, samples were divided into their Day 0 post-stress bleaching severity groups (low, moderate, and high). Points with no further lines represent mortality, and point represents a coral nubbin on a given day. Colored lines highlight the 50% most frequent trajectories, whereas gray lines are low support trajectories. Each plot includes three boxes to show the bleaching severity groups: blue = low, yellow = moderate, and red = high. Plots were made using otsplot within the R (version 4.0.5) package vcrpart.

**Figure 3.**
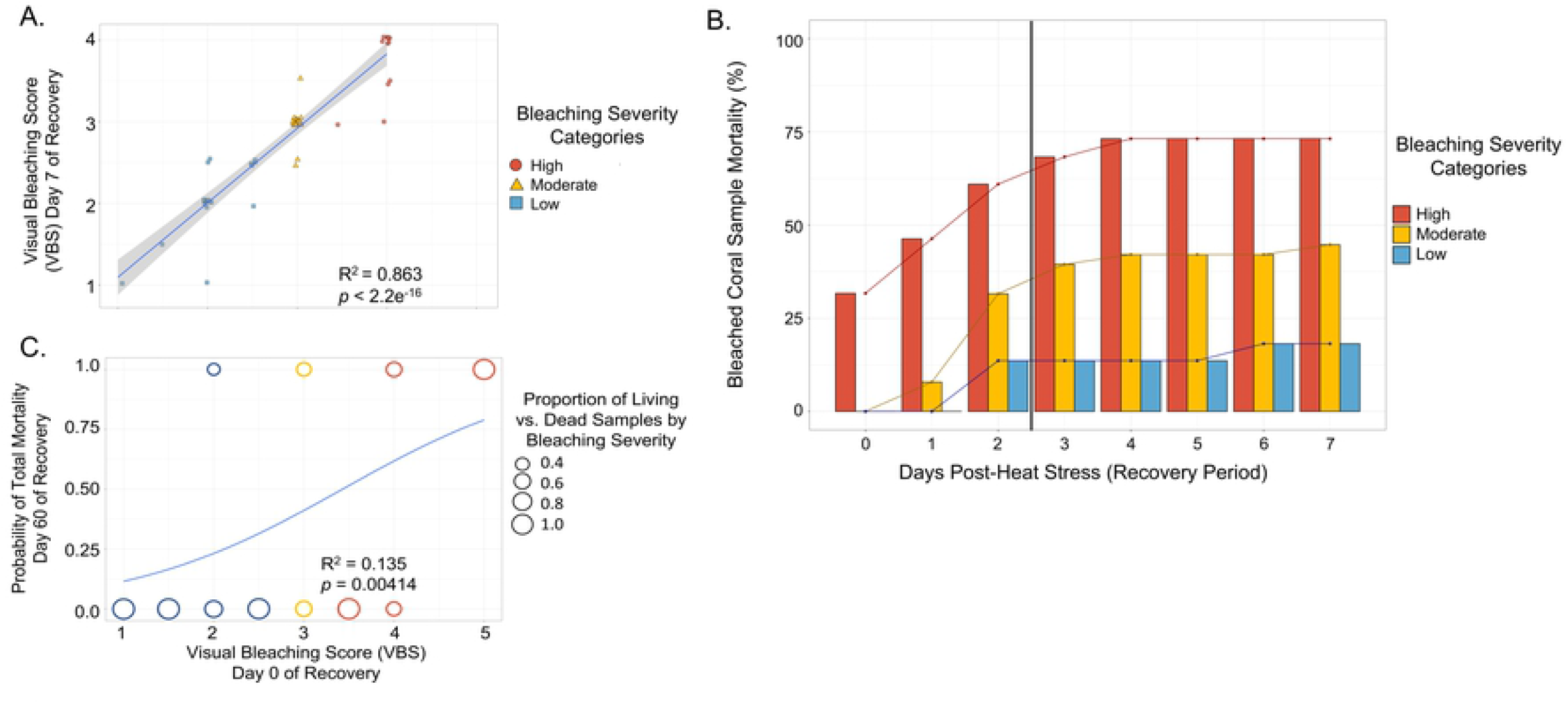
Predictability of bleaching severity immediately after heat stress (Day 0 post-stress) for survival. (A) Scatterplot showcasing sample bleaching severity on Day 0 versus Day 7 post-stress (based on visual bleaching score, VBS, and with bleaching severity categories), including a line of best fit with 95% confidence intervals (linear regression). A linear mixed effects model was used, with reef region as a random intercept to account for spatial variability, and the marginal R^2^ value is provided. (B) Percentages, per bleaching severity group, of heat stressed corals with total tissue mortality daily from Day 0 to Day 7 post-stress. Mortality during the heat stress experiment (n=13) was attributed to the high bleaching severity category (i.e., low heat resistance), and severely bleached coral nubbins that died throughout the post-stress week were added to this count. A connected scatterplot of each bleaching severity group’s mortality over time is included, as well as a black vertical line to indicate the day on which 50% of all observed mortality was surpassed. (C) Binomial representation of living (value = 0) and dead (value = 1) nubbins 60 days after heat stress versus bleaching severity on Day 0 post-stress, where circle sizes represent proportion of living vs. dead samples at each VBS category (mixed effects logistic regression, accounting for spatial variability, and the pseudo R^2^ value is provided). Circle colors correspond to bleaching severity immediately after heat stress: blue = low bleaching, yellow = moderate bleaching, and red = high bleaching.

Mortality post-stress was also predictable after 16 hours. By the end of one-week post-heat stress, 51 total nubbins had died (50.5% mortality), including 38 (43.2%) among the eighty-eight nubbins that survived immediate impacts from the heat stress experiment. Further, more than 50% of total mortality occurred within two days of the heat stress experiments. Coral nubbins with little to no bleaching on Day 0 survived best: only 4 out of 22 (18%) nubbins in this category died, three of them by Day 2. Nubbins showing moderate bleaching had higher mortality: 17 out of 38 nubbins died (45%), 12 of these by Day 2. The most heat susceptible nubbins experienced the highest mortality: 30 out of 41 (73%, including those that sustained severe to total bleaching or died during heat stress) (Fig. 3B). Overall, bleaching severity on Day 0 post-stress was significantly and positively correlated with likelihood of mortality by Day 7 (Fig. S3A, mixed effects logistic regression, *p* = 0.00184, pseudo marginal R^2^ = 0.165) and Day 60 (Fig. 3C, logistic regression, *p* = 0.00414, pseudo marginal R^2^ = 0.135), although relatively low R^2^ values show that much of the variation remained unexplained. Overall, bleaching severity on Day 0 post-stress was not as predictive of mortality at Day 7 as were parallel data for bleaching severity throughout the weeklong period (see above).

### Environmental differences in bleaching resistance

Previous work in Palau (Cornwell et al., 2021) showed widespread occurrence of heat resistance, though presence of heat resistant corals in part coincides with thermal environment. Consistent with Cornwell et al., (2021), we found that bleaching resistance was widespread throughout Palau’s geographic regions. There were minimally bleached coral nubbins that originated from 11 patch reefs (7 in the south and 4 in the north) and three fore reef locations (1 in the south and 2 in the north). Four of these patch reefs (2 in the south and 2 in the north) had all categories: low, moderate, and high bleaching severity. Coral nubbins that sustained exclusively moderate or worse bleaching originated from ten of 21 patch reefs, a slightly lower proportion than the fore reef sites, where four (of seven) exhibited only moderate or worse bleaching (Fig. 4A).

**Figure 4.**
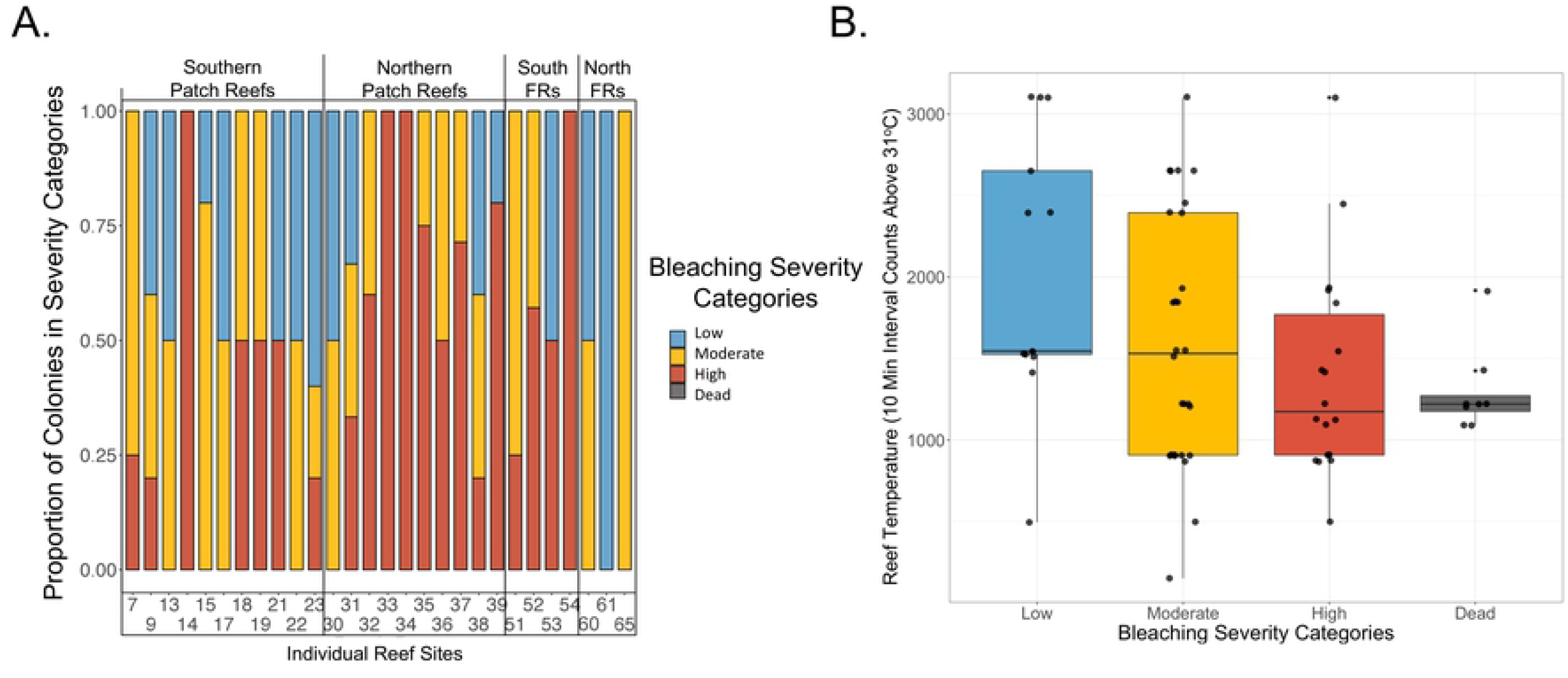
Bleaching severity across variable reef environments and thermal regimes. (A) Stacked barplot showing proportion of bleaching severity categories (low, moderate, and high bleaching) across the twenty-eight sampled reef sites. Reefs are organized by the following general groups: southern and northern patch reefs, and southern and northern fore reefs (written as FRs). (B) Boxplot showing the relationship between reef temperature extremes (represented by 10 minute interval HOBO logger counts above 31°C) and bleaching severity groups (low, moderate, and high bleaching). An ordinal logistic regression was performed on increasing bleaching severity groups (here from low severity to dead), whereby only the low and moderate bleaching severity groups differed significantly (*p* < 0.05).

Cornwell et al., (2021) related bleaching differences on Palauan reefs to their respective temperature regimes, so we used our independent data to assess whether temperature extremes (i.e., recorded 10-minute interval events above 31°C) of the originating reefs predicted how corals were able to survive heat stress. We found a weak negative relationship between the three bleaching severity categories (Low, Moderate, and High) immediately after heat stress and reef temperature, in which only the low and moderate bleaching severity categories significantly differed (ordinal logistic regression, *p* < 0.0001). Samples that died during the heat stress experiment also tended to originate from cooler reefs, though further study is needed to address whether reef temperature strongly influences coral heat stress responses in this system (Fig. 4B). However, there was also a large amount of variation in the reef specific data. For example, the four reef sites that had only severely bleached coral nubbins ranged in temperature extremes from well below (495.5, PR 33) to above the reef mean counts above 31°C (2652.3, PR 14).

Similarly, there was one reef that had entirely minimally bleached coral nubbins (Northern fore reef site 61) but a moderate count of temperature extremes (1526) (Table S1). This suggested a general link between reef temperature and heat stress resistance but also highlighted that there are likely other environmental factors influencing resistance.

### Health of coral nubbins beyond one week post bleaching

Because visual bleaching recovery did not occur in samples over the first week, we returned at one- and two-months post-stress to evaluate bleaching and mortality in the remaining samples. Out of the fifty surviving coral nubbins, thirty-seven achieved full visual recovery (VBS 1, 74%) after approximately one month. The remaining corals also all fell under the low bleaching severity category with minimal observed bleaching. There were representatives from all three original bleaching severity categories among those fragments (Fig. 5). After approximately two months, forty-nine out of fifty heat stressed nubbins were still alive, and all had recovered fully (VBS 1) (Fig. 4). All nubbins had a high probability of surviving and visibly recovering by two months post-bleaching if they reached Day 7 of the post-heat stress period. These collective results showed that bleaching sustained during short-term heat stress did play a significant role in the likelihood of survival past one week, though there are likely other important factors apart from bleaching severity to consider (Fig. 3C, Fig. S3, Fig. 5).

**Figure 5.**
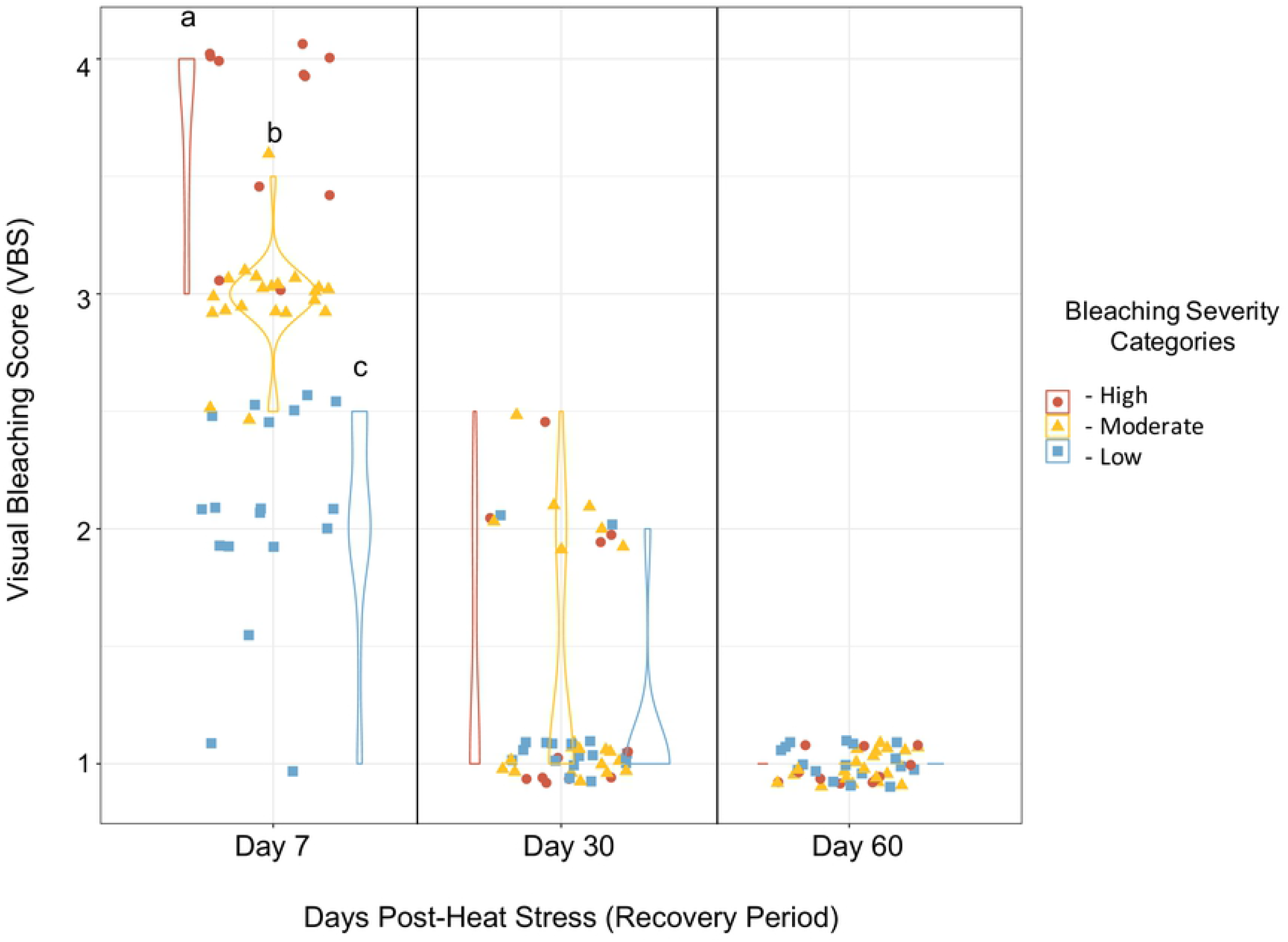
Visual bleaching recovery in corals beyond one week after heat stress. Each point per panel denotes a coral nubbin (representing an individual coral colony), nubbins that died by Day 7 of the recovery period (n=51) were excluded. Bleaching severity was measured using visual bleaching scores (VBS) at approximately Day 30 and Day 60; see Table S1 for the complete list of exact days for each nubbin. Note that there were a few samples with in between VBS scores (designated by increments of 0.5), for which the two observers could not decide on a single bleaching category. Tukey tests were performed on groups at each timepoint, with a significant threshold of *p* ≤ 0.05. Bleaching severity between all resistance groups on Day 7 post-stress differed significantly where *p* < 0.00001, but not on Days 30 and 60.

## Discussion

We tested how *Acropora hyacinthus* coral nubbins respond to heat stress in commonly used short-term heat stress experiments that are normally assayed within 16 hours post-stress (Grottoli et al., 2021). Pigmentation loss in nubbins was highly stable under lab observation from 16 hours to seven days post-stress. In addition, coral nubbins that survived three days after heating had high survival and recovery after 1-2 months in the lab. These data confirm that bleaching is consistently stable days after a short-term heat stress experiment concludes and begin the process of ground-truthing whether short-term bleaching experiments can predict longer term survival.

This experimental design also allows for testing links between variation in heat resistance and recovery. Further, we confirmed widespread variation in heat resistance and recoverability in variable thermal regimes and environments throughout Palau (see also Cornwell et al., 2021).

### Reliability and efficiency of a non-destructive visual bleaching score metric

We utilized a five-point visual bleaching scoring metric—(1) none (2) visible, (3) moderate, (4) severe, and (5) total bleaching—in order to quickly and effectively evaluate a large sample size of control and heat stressed coral nubbins. Visual bleaching scores were advantageously nondestructive and nondisruptive, meaning we did not sacrifice any portion of tracked colony fragments or disturb fragments to assess bleaching during the post-stress period. Several studies have previously relied solely on qualitative visual scoring methods (e.g., Mills et al., 2013; Thurber et al., 2014; Gintert et al., 2018; Fisch et al., 2019), while others have calibrated visual results with quantitative assays like symbiont concentration via flow cytometry or sequencing and chlorophyll concentration (e.g., Siebeck et al., 2006; Bay et al., 2016; Thomas and Palumbi 2017; Fuller et al., 2020). Similarly, we confirmed our visual bleaching scores with flow cytometry data using nubbin replicates sacrificed after the heat stress experiment. We found that flow cytometry results matched well with the three main bleaching severity categories established with visual scoring (none to visible bleaching, i.e., high resistant; moderate bleaching, i.e., moderately resistant; and severe to total bleaching, i.e., low resistant). However, we were unable to distinguish between some individual categories along the 5-point scale: none vs. visible and severe vs. total categories had similar symbiont concentrations. Previous studies have shown that pigmentation loss may correlate with chlorophyll and/or symbiont concentration depending on the individual (Jones 1997) or species (Rodrigues and Grottoli 2007; McLachlan et al., 2021; Chapron et al., 2022), and healthy coral colonies can also have highly variable photosynthetic pigment concentrations (Apprill 2007). Therefore, starting pigmentation or chlorophyll vs. symbiont concentration differences may have influenced quantitative correlation with qualitative visible bleaching in these *Acropora hyacinthus* colonies. Overall, confirming our qualitative results with a quantitative method allowed us to go forward with broad bleaching severity categories and showcased the importance of combining methodologies when evaluating bleaching. Together, this validated the merit of employing a reliable visual scoring metric in place of more expensive and time-consuming methods such as flow cytometry when performing experiments that require large sample sizes.

### Post bleaching trajectories are highly stable among corals with variable bleaching resistance

We found high intraspecific variation in bleaching sensitivity after the short-term heat stress experiment, comparable to other studies in which colonies were exposed to high temperatures at similar and variable lengths of time (Morikawa and Palumbi 2019; Kavousi et al., 2020; Cornwell et al., 2021). One week after heat stress, visual bleaching scores for coral nubbins were not significantly different than they were after 16 hours (Fig. 2). Among all corals that survived the first week after heat stress, 90% (45 nubbins) remained in the same bleaching severity category and 4 out of 5 of the other corals improved bleaching categories. The lack of further visible symbiosis breakdown after heat stress suggested that symbiont loss was mostly restricted to the discrete heat stress event, while any remaining symbionts maintained an association with their host during the seven-day post-heat stress period. However, these remaining symbionts’ systems such as photosynthetic ability and translocation of energy to the host may be significantly impaired during early holobiont stress recovery (Takahashi et al., 2008; Gierz et al., 2016; Gierz et al., 2017).

Mortality also followed these patterns. We found that 90% of mortality occurred within the first few days after short-term heat-stress. Mortality was high among corals with high bleaching within the first 3 days of heat stress (56%) but was low thereafter (another 7%). Mortality was far lower for nubbins with low or moderate bleaching (30%) and dropped to near zero after Day 3 (Fig. 2). Higher mortality in corals with severe bleaching is consistent with what has been shown in a recent long-term study (Matsuda et al., 2020).

### Predictability of survival beyond one week after heat stress

We recorded a total mortality rate two months after short-term heat stress (∼51%) that was comparable to other studies on bleaching susceptible species (e.g., 60% in *Montipora capitata* Rodrigues and Grottoli 2007; see also Baird and Marshall 2002; Williams et al., 2017). However, our serial monitoring of time points showed that mortality visible after two months actually occurred within 3 days of bleaching for our system. One month after heat stress, we recorded that all coral nubbins were in the low bleaching severity category and the majority were fully recovered. Corals of all resistance types were among those that still had some visible bleaching after one month. We found high survival and full recovery among all nubbins two months after heat stress, even for those that had been severely bleached.

These data also show a significant correlation between survival after one week and likelihood of visible recovery in the future, which suggests that the first week after bleaching may be a critical period for short-term heat stressed corals. Indeed, most of the corals that were scored as severely or totally bleached died within 3 days of the heat stress experiments. By contrast, virtually all the corals that were only minimally bleached in our stress tests survived. Similar results have been shown in field surveys of corals days to months after natural bleaching events (e.g., Jones 2008; Bahr et al., 2017; DeCarlo et al., 2017) and report that severely bleached, white colonies seldom survive. There was variation in the two month post-stress mortality results that could not be explained by early bleaching status. Therefore, further research into other potential factors, such as prevalence of diseases (Muller et al., 2108; Walton et al., 2018; Brodnicke et al., 2019) and access to nutrients (Wooldridge and Done, 2009; Burkepile et al., 2019; Morris et al., 2019) would be beneficial.

### Intraspecific variation in thermal resistance and recovery throughout Palau

We found that high bleaching resistance was geographically widespread across variable environments though more common in corals coming from hotter reefs, with similar results reported in Cornwell et al., (2021). A minority of reefs had corals of all bleaching resistant types. We report that half of all reefs had highly resistant corals, while 71% of all reefs each had low resistant and/or moderate resistant corals. Furthermore, reef temperature extremes were weakly associated with bleaching severity (e.g., Oliver and Palumbi 2011; Barshis et al., 2013), and with post-stress mortality (e.g., Pineda et al., 2013).

We also report for the first time that recovery from severe bleaching may also be a widespread trait for *Acropora hyacinthus* in Palau: coral nubbins from all geographic regions survived and had visible recovery in our experimental system. We recorded 100% mortality of bleached coral nubbins from only six out of the 28 reef sites. This suggests that corals with high bleaching recoverability may be located throughout Palau. It is also possible that other facets of environmental variation could similarly impact heat resistance and recovery. For example, intensity and duration of variability, accumulated heat stress (e.g., degree heating weeks), or longer-term oscillations like El Niños may play a more consequential role predicting the likelihood of heat resilience and survival (McClanahan et al., 2007).

### Implications and future directions

Short-term heat stress experimental designs (McLachlan et al., 2020) have been employed as a powerful tool to globally and rapidly assess susceptibility to bleaching and likelihood of survival in susceptible corals on a variety of reef types. An advantage of this approach is the ability to test many coral nubbins rapidly within 2-3 days of collection (Palumbi et al., 2014; Seneca and Palumbi, 2015; Traylor-Knowles et al., 2017; Thomas et al., 2018). This minimizes experimental effects such as acclimation to lab conditions and reduces the chance of other sources of mortality or stress such as starvation. However, these advantages must be weighed against the fact that minimal post-collection recovery time could confound transportation and heat stress, and the strong heat pulse could impair coral function so quickly that it might take days or longer for the full reaction to become apparent (Grottoli et al., 2021). In our dataset, non-stressed control nubbins were highly robust, such delayed reactions among heat stressed nubbins were rare, and rapid assays of bleaching were accurate measures of longer-term response. Physiological metrics such as chlorophyll density, Symbiodiniaceae density, and host protein content can differ in their ability to resolve temperature and site-specific differences between short-term (18 hours) and long-term (21 days) heating assays (Voolstra et al., 2020). Other potential disadvantages are that standardized experimental heat stress applied to nubbins from variable reef environments may not fully capture the natural variability these corals experience nor the spectrum of natural bleaching events that are known to vary in duration and intensity. Therefore, it is imperative to combine reliable standardized protocols, which can reveal initial integral mechanistic patterns, with studies that integrate other environmental factors and variable durations and intensities of heat stress.

This study was conducted solely on the bleaching sensitive species *Acropora hyacinthus*. Species in this genus react quickly to strong heat stress, whereas other genera such as *Pocillopora* (e.g., D’Croz and Maté 2004; Poquita-Du et al., 2020) or *Porites* (Kenkel et al., 2013) may require longer exposure to lower levels of heat to bleach without immediate death. One important future direction is to use this experimental design on other prominent reef-building coral species and in other variable reef types. Future assays should test other possible environmental (e.g., currents and water flow, more extreme temperature differences, available nutrients or predators, and light), physiological (metabolic baseline, energy reserves and heterotrophy, symbiont temperature resilience), and genomic predictors of thermal resilience during this short time period—as these factors are known to relate to thermal sensitivity over longer timescales (e.g., McClanahan et al., 2005; Borell et al., 2008; Schoepf et al., 2015; Levas et al., 2018; Thomas et al., 2019; DeCarlo et al., 2020; Rosic et al., 2020; Schoepf et al., 2020; Chapron et al., 2022).

### Conclusions

We have used a simple, rapid, and low-cost experimental design to test coral nubbins for bleaching resistance and recovery. This coupling is important, because resilience against ocean warming may require a combination of resistance and recovery (Levin and Lubchenco, 2008; Bernhardt and Leslie, 2013). Our data confirm that bleaching and mortality recorded quickly after short-term heat stress experiments are stable and reliable measures of the stress reaction. In addition, our findings that high heat resistance and recoverability may be widespread throughout Palau and that low resistant coral nubbins can also recover well suggest that many *Acropora hyacinthus* individuals may have the metabolic machinery necessary to effectively resist and/or recover from heat stress. Ultimately, prioritizing the highest bleaching resistant corals may help maintain resilience on reefs in the future (van Oppen and Gates 2006), but including corals of variable resistance and recoverability in reef management plans may further increase diversity and sustainability.

## Author Contributions

Conceptualization: N.S.W. and S.R.P. designed the experiment.; Data Curation: N.S.W., S.R.P., B.H.C., K.C.A., and V.N. collected coral samples for the experiment. N.S.W. conducted the experiment and collected data under the supervision of S.R.P., and V.N. aided in experiment data collection.; Formal Analysis: N.S.W.; Writing, Original Draft: N.S.W.; Writing, Review & Editing: N.S.W., S.R.P., B.H.C., K.C.A., V.N., and Y.G.; Funding Acquisition: S.R.P. and Y.G.

## Acknowledgements

We thank the staff and boat operators at the Palau International Coral Reef Center, and Stanford University undergraduate interns Callan Hoskins, Colin Hyatt, Mehr Kumar, and Julien Ueda for their assistance collecting samples and running experiments in the field. We also acknowledge current and former members of the Palumbi lab—Elora López-Nandam, Erik Hanson, Kristin Robinson, and Marilla Lippert—as well as Andrea G. Grottoli, Elizabeth Hadly, John R. Pringle, and referees during the review process for their comments and edits on the manuscript. This study was supported by NSF grant OCE [1736736], and by support through Stanford University’s office of Development.

## Supplementary Figures and Data Captions

**Figure S1. Geographic locations of all reefs and reef groups**. Map of Palau with relative reef positions, created using ArcGis. Reefs are circled based on reef cluster groups; patch reefs are circled in red, yellow, green, and blue, and fore reef sites are circled in purple. Latitudinal and longitudinal coordinates of all reefs are included and written in decimal degrees.

**Figure S2. Bleaching resistance variation following heat stress**. Counts of control (blue) and bleached (green) colony samples before and after the two-day short-term heat stress experiment. Bleaching severity before and after heat stress was measured by visual bleaching score (VBS).

**Figure S3. Relationship between bleaching severity immediately after heat stress (Day 0 post-stress) and mortality on Days 7 and 30 post-stress**. Binomial representation of living (value = 0) and dead (value = 1) coral nubbins 7 days (A) and 30 days (B) after heat stress versus bleaching severity on Day 0 post-stress, where circle sizes represent proportion of living vs. dead samples at each VBS category (mixed effects logistic regression, accounting for spatial variability, with pseudo R^2^ values provided). Circle colors correspond to bleaching severity immediately after heat stress: blue = low bleaching, yellow = moderate bleaching, and red = high bleaching.

**Figure S4. Breakdown of bleaching severity across geographic locations and reef temperatures**. (A) Violin plot of bleaching severity and geographic locations. Bleaching severity is measured through visual bleaching scores (VBS 1, no bleaching, to VBS 5, total bleaching), and each point denotes a nubbin that represents a coral colony within geographic locations. All locations are in lagoons apart from the Fore Reef category, and colors correspond to groups in Figure S1. We ran an ANOVA and Tukey test for significance (Table S2), * denotes *p* ≤ 0.05. (B) Scatterplot showing the relationship between reef temperature extremes (represented by 10-minute interval HOBO logger counts above 31°C) and visual bleaching score with 95% confidence intervals. Day 0 and 7 after heat stress are shown. Linear regression results: Day 0, *p* = 0.02167, R^2^ = 0.0731, df = 57, and Day 7, *p* = 0.0292, R^2^ = 0.1287, df = 28.

**Figure S5. Photographic nubbin representations of the 5 visual bleaching score categories and total mortality**. Two observers determined each nubbin’s visual bleaching score category and mortality. Here, one nubbin example is provided per visual bleaching score category and mortality category.

**Table S1. Metadata spreadsheet**. Visual bleaching scores, symbiont proportion, temperature data, and descriptions of reef and resistance/bleaching severity groupings.

**Table S2. Summary of statistics results**. All statistics result outputs, including linear regressions, logistic regressions, ANOVAs, and post-hoc Tukey tests.

